# Knockdown of *NeuroD2* leads to seizure-like behaviour, brain neuronal hyperactivity and a leaky blood-brain barrier in a *Xenopus laevis* tadpole model of DEE75

**DOI:** 10.1101/2023.12.06.570491

**Authors:** S. Banerjee, P. Szyszka, C.W. Beck

**Affiliations:** Department of Zoology, University of Otago, PO Box56, Dunedin, New Zealand; Brain Health Research Centre, University of Otago, Dunedin, New Zealand; Genetics Otago Research Centre, University of Otago, Dunedin, New Zealand

**Author notes:** Correspondance to: Caroline Beck.

**Keywords:** Epilepsy, NeuroD2, blood-brain barrier, Losartan, developmental and epileptic encephalopathy, C-starts, Mauthner neuron

## Abstract

Developmental and Epileptic Encephalopathies (DEE) are a genetically diverse group of severe, early onset seizure disorders. DEE are normally identified clinically in the first six months of life by the presence of frequent, difficult to control seizures and accompanying stalling or regression of development. DEE75 results from *de novo* mutations of the *NEUROD2* gene that result in loss of activity of the encoded transcription factor, and the seizure phenotype was shown to be recapitulated in *Xenopus tropicalis* tadpoles. We used CRISPR/Cas9 to make a DEE75 model in *Xenopus laevis*, to further investigate the developmental aetiology. *NeuroD2*.*S* CRISPR/Cas9 edited tadpoles were more active, swam faster on average, and had more unprovoked escape responses (C-starts) than their sibling controls. Live imaging of Ca^2+^ signalling revealed prolongued, strong signals sweeping through the brain, indicative of neuronal hyperactivity. While the resulting tadpole brain appeared grossly normal, the blood-brain barrier was found to be leakier than that of controls. Additionally, the TGFβ antagonist Losartan was shown to have a short-term protective effect, reducing neuronal hyperactivity and reducing permeability of the blood-brain barrier. Severity of the behavioral phenotype correlated with increased with editing efficiency. Our results support a haploinsufficiency model of DEE75 resulting from reduced NeuroD2 activity during vertebrate brain development, and indicate that a leaky blood-brain barrier contributes to epileptogenesis.

## Introduction

Epilepsy is one of the most common neurological disorders, estimated to affect 1% of the population at some time in their lives. Seizures, the defining feature of epilepsy, are visible manifestations of the underlying hyper-exictability of neuronal firing in the brain (Gross 1992). Patients are generally treated with one or more anti-seizure drugs (ASD) which aim to reduce or eliminate seizures. However in many patients, seizure control cannot be achieved and the ongoing and unpredictable nature of seizure occurance can result in poor quality of life. The developmental and epileptic encephalopathies (DEE), which present in infancy or very early childhood, are the most severe group of genetic epilepsies (Mctague *et al.* 2016). DEE incorporates previously described syndromes of infantile onset epilepsy such as Otahara, Dravet and West syndromes, as well as later onset syndromes such as Lennox Gastaut (reviewed in (Scheffer and liao 2020). Diagnosis is confirmed by the presence of seizures, abnormal electroencephalogram (EEG) activity that persists between seizures, and stalling, delay of regression of cognative and behavioural development (Scheffer *et al.* 2017; Scheffer and liao 2020). Mortality is 25% by age 20 (Camfield and camfield 2008), and survivors often have significant lifelong intellectual and motor disabilities, requiring lifelong care (Keezer *et al.* 2016).

DEE are all linked by a similar clinical presentation: early onset, unrelenting seizures that are refractory to treatment. Individual DEE are ultra-rare, but the estimated cumulative incidence of DEE is 169 in 100,000 (Poke *et al.* 2023). Genome and whole exome sequencing of patients with early onset epilepsy e.g. (EPI4K *et al.* 2013; Epi 2016), has identified 110 genes with rare variants that cause DEE, curated in Online Mendelian Inheritance in Man database (OMIM). However, this may just be the tip of the iceberg, as a recent review manually curated 936 genes associated with monogenic epilepsy, implicating 4% of human protein coding genes (Oliver *et al.* 2023). Of these, 818 (89%) were DEE-associated, with around 30% autosomal dominant (AD) inheritance (likely *de novo*) 60% autosomal recessive (AR) with the remaining 10% X-linked or mitochondrial (Oliver *et al.* 2023).

Functional studies in animal or cell models are lacking for the majority of DEE, and with so many genes and variants implicated, new approaches are needed to manage these severe conditions. Recent work demonstrates that the tadpole is a good model for human brain development (Willsey *et al.* 2021; Ta *et al.* 2022) as vertebrate central nervous system (CNS) development follows remarkably conserved mechanisms (reviewed in (Exner And willsey 2021). In fact, much of what we know of vertebrate CNS development was discovered from studies in *Xenopus* and other amphibian embryos. The large size, plentiful number and external development of *Xenopus* eggs makes them amenable to genetic manipulation and has led to accurate fate maps allowing this to be targeted. Furthermore, early functional studies in *Xenopus* led to the current understanding of neural induction, patterning and neurogenesis in vertebrates (reviewed in (Exner And willsey 2021). The week-old *Xenopus* tadpole brain is organised similarly to the human brain, with the main difference being the expansion of forebrain into the six-layered neocortex of humans, which is formed from the inside out via a process of radial migration, and generating the sulci and gyri (Molnar *et al.* 2006; PARIDAEN AND HUTTNER 2014). *Xenopus* forebrains are lissencephalic (smooth), unlayered, and form from the outside in, so the oldest neurons are found on the outside (Moreno And gonzalez 2017). While higher cognitive behaviours cannot be studied in *Xenopus*, the tractability of the tadpole system, combined with direct access to the brain at all developmental stages, make it a powerful model system for investigating the origins of human disorders arising from aberrant brain development (PRATT AND KHAKHALIN 2013). Recent examples where *Xenopus* tadpoles have been used as models of human brain disorders include autism (Willsey *et al.* 2020; Willsey *et al.* 2021), fragile X syndrome (Truszkowski *et al.* 2016), and DEE (Yoo *et al.* 2017; Sega *et al.* 2019).

Here, we set out to build on previous work by Sega et al (Sega *et al.* 2019), who used *X. tropicalis* tadpoles to show that two *de novo* pathogenic variants in *NEUROD2* cause DEE75. *Xenopus laevis* tadpoles have a well developed brain at Nieuwkoop and Faber (NF) stage 47, which is about one week of age, and pre-feeding. Seizures can be chemically induced by adding pentyleltetrazole (PTZ) or 4-aminopyrolidine (4-AP) to the swimming medium (Hewapathirane *et al.* 2008; Bell *et al.* 2011; Panthi *et al.* 2023). Additionally, since the skull has yet to form at this stage, the brain is easily accessible for electrode recording of local field potentials or Ca^2+^ signalling (Hewapathirane *et al.* 2008). The tadpoles can be housed individually in wells of a 24-well culture plate with 1 ml of medium, for automated behavioral classification and drug testing (Panthi *et al.* 2023). We wanted to see whether automated quantification of tadpole behaviour would be sensitive enough to enable drug testing for anti seizure activity in this model. We also showed that brain hyperactivity, characteristic of spontaneous seizures, can be detected in live tadpole brains using the genetically encoded Ca^2+^ sensor GCaMP6s. While brain morphology appeared unaltered in CRISPants, the blood-brain barrier (BBB) was less able to prevent the loss of sodium fluorecein (NaF) dye injected into the ventricle, suggesting that a leaky blood brain barrier may contribute to the epileptogenic brain of DEE75 children. Our findings show that *X. laevis* tadpoles are a useful model of DEE75, and may be useful in future to develop models of other DEE for functional study. These models could be used to inform and undertake pre-clinical testing of re-purposed drugs, such as Losartan, which may have the potential to protect the brain from the damaging effects of unrelenting seizures, thereby improving the quality of life for patients.

## Materials and methods

### Production of *Xenopus laevis* eggs and embryos

Animal procedures were approved by the University of Otago’s Animal Ethics Committee under animal use procedures AUP19/01 and AUP22/12. Adult female *Xenopus laevis* were induced to lay eggs by injecting with 500 IU per 75 g body weight of Human chorionic gonadotrophin (Chorulon) into the dorsal lymph sac, 16 hours before eggs were required. They were placed in a dark incubator overnight in pairs in small holding tanks containing “frog water” (tap water passed through carbon filters to remove chlorine). After egg laying commenced, each female was placed in 1 L of 1 x Marc’s modified ringers (MMR: 10 mM NaCl, 0.2 mM KCl, 0.1 mM MgSO_4_.6H_2_0, 0.2 mM CaCl_2_, 0.5 mM HEPES, 10 μM EDTA, pH 7.8) and eggs collected hourly and fertilised using fresh testes from a euthanised adult *X. laevis* male. Jelly coats were removed immediately following embryo rotation (15-20 minutes), using a 2% solution of Cysteine HCl (pH 7.9) and the resulting de-jellied embryos rinsed three times in MMR. Embryos were raised in small batches in 0.1 x MMR with no antibiotics and staged according to Nieuwkoop and Faber’s staging series (Nieuwkoop And faber 1967).

### CRISPR/Cas9 targeting of *NeuroD2*

ChopChop v2 (Labun *et al.* 2016) was used to design short guide RNA (sgRNA). As *X. laevis* is allotetraploid (Session *et al.* 2016; Fisher *et al.* 2023), the sgRNA was designed to the *S* form, which is more highly expressed in tadpole brains (Ta *et al.* 2022). SgRNA were chosen based on specificity and ability to edit both *L* and *S* homeologues (with no other chromosomal off-targets). sgRNAs rnk5 and rnk20 (ranked 5th and 20th, respectively, for efficiency by ChopChop) met these criteria and were also located near the DEE75 human variant sites. Editing efficiency was predicted using InDelphi (Shen *et al.* 2018) (https://indelphi.giffordlab.mit.edu/), (Figure S1) which is a good editing predictor for *Xenopus* (Naert *et al.* 2020). *NeuroD2* sequence for human and *X. laevis* L and S proteins and *X. laevis NeuroD2*.*S* mRNA were obtained from NCBI and imported into SnapGene v5.2.4. Protein sequences were aligned in SnapGene using MUSCLE alignments and the bHLH, NLS domains and human DEE-causing variants annotated from the human sequence and data in previously published studies (Sega *et al.* 2019; Mis *et al.* 2021). Using the InDelphi (Shen *et al.* 2018) online prediction tool, 60 bp up- and downstream of the target Cas9 site in *X. laevis NeuroD2*.S were entered in “single mode” using mESC cell type, for each sgRNA. The EnGen sgRNA template oligo designer tool (NEB) was used to design long 54-55 bp oligonucleotides for sgRNA synthesis, with the 20 nucleotide sequence from ChopChop v.2 (without the “NGG” PAM). A “G” is added to the 5’ end of the target sequence if not present, as this enables optimal transcription. Primer and sgRNA oligonucleotide sequences can be found in Table 1, predicted edits for each sgRNA can be found in Figure S1.

**Table 1.**
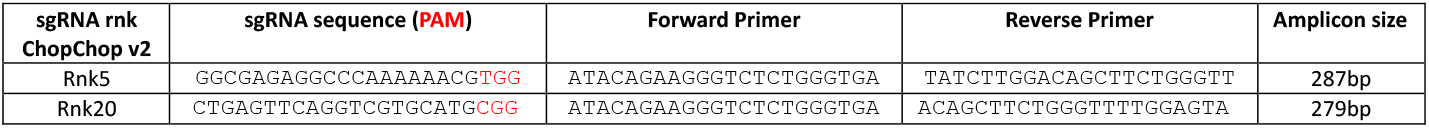
sgRNA sequences and amplification/sequencing primers for this study.

Oligonucleotides were ordered from Integrated DNA Technologies (IDT), and used with the EnGen sgRNA synthesis kit, *S. pyogenes* (NEB) to make sgRNA. The resulting sgRNA was dissolved in water, stored as small aliquots at -80°C and thawed on ice immediately prior to use. 0.3 μl of EnGen *S*.*pyogenes* Cas9 NLS enzyme (protein) was added to sgRNA, resulting in a 3 to 5 fold dilution, and incubated at 37°C for 5 minutes. The resulting mixture was loaded by back-filling into a pulled Drummond glass capillary needle. Fertilised, de-jellied embryos within 1 hour of fertilisation were checked for sperm entry points and lined up in a row in a 2 mm x 40 mm trench cut into a 50 mm petri dish lined with 2.4 ml of 2% agar and filled with 6% Ficoll PM400 in MMR and placed under a dissecting microscope for microinjection. 9.2 nl of Cas9/sgRNA were injected into each egg near, but not into, the female pronucleus, using a Drummond Nanoject II injector held with a Narishige MM3 micromanipulator. Total amount of sgRNA per egg was 400 pg for the rnk5 and 900 pg for the rnk20 sgRNA. Eggs were injected in batches of 25 and immediately placed into the incubator at 18°C following injection. Each concentration of sgRNA was injected into 100 embryos from the same sibship, and controls were generated using only Cas9 enzyme (no sgRNA). After 2-3 hours, the embryos were checked for normal development and moved to 3% Ficoll in 0.1 x MMR. The following day embryos were checked for normal development, and five embryos were chosen at random for genotyping. The remainder were moved to 0.1 x MMR until stage 46-47 was reached. Genotyping was done by homogenising each embryo or tadpole in 200 μl 5% Chelex beads in phosphate buffered saline with 1.5 μl Proteinase K (25 mg/ml) and incubating for 3 hours (embryos) to overnight (tadpole) at 65°C. The reaction was then terminated by incubating samples at 95°C for 5-10 minutes, and briefly centrifuged to pellet the beads. 1 μl of the supernatent was used directly for PCR (for primers, see Table 1). TIDE analysis (Brinkman *et al.* 2014) (https://tide.nki.nl/) of CRISPant samples vs. controls, which only received Cas9 protein, was used to confirm editing and InDelphi predictions.

### Recording and analysis of stage 47 tadpole seizure behaviour

Individual tadpoles were transferred to a 24 well culture plate with each well containing 1 ml of 0.1 x MMR, enough to submerge tadpoles, allowing free swimming. A Panasonic DMC- FZ1000 camera was mounted on a tripod above the tadpoles and the Movie mode was used to capture 30 minutes of MP4 video at 25 frames per second. Tadpoles were backlit with an array of 176 ultrabright LED lights (Neewer, 1300 lumens). In TopScan, the locomotion super module was used to create 24 arenas, and a clear background. Swimming tracks were generated for each tadpole, representing 30 minutes of activity. Mean velocity (mm/sec) was calculated by TopScan for each tadpole as in (Panthi *et al.* 2023). C-shaped contractions (C-SC) over a 10-minute period were determined by the software counting the total number of C-SC events that met both the elongation ratio (<1.5) and motion measure (>0.2) criteria, and confirmed by manual checks. Examples of postures scored as C-SC in video frames can be found in Figures 1B and 2B, with a positive score if the tadpole’s head and tail described more than half of a complete circle. C-SC were also grouped into clusters, where multiple C-SCs frames were detected with an interval of less than 2 seconds between them.

**Figure 1:**
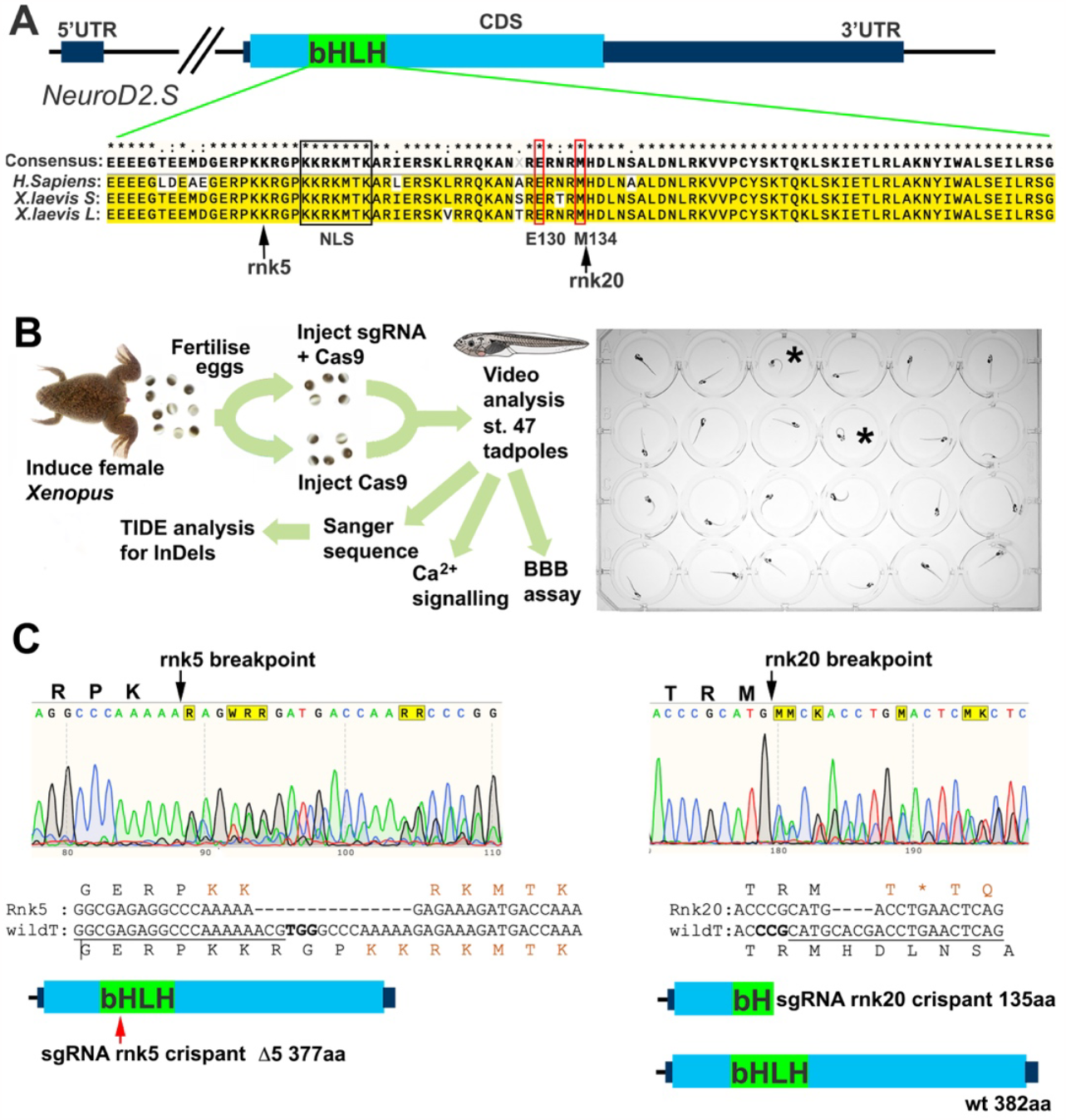
CRISPR/Cas9 edi4ng of NeuroD2 in tadpoles. **A)** Schematic of the *X. laevis NeuroD2*.*S* gene, the homeologue *NeuroD2*.*L* is the same for all key aspects shown, including sgRNA binding sites. The NeuroD2 transcription factor is encoded by a single exon (CDS) and the bHLH DNA binding domain is indicated in green. Alignment of the bHLH domain between *H. sapiens* and *X. laevis* shows a high degree of conservation at protein level. Nuclear localisation sequence (NLS) is indicated by a black box, and the conserved positions of the previously described *de novo* human variants E130Q and M134T that cause DEE75 (Sega *et al.* 2019) indicated by red boxes. The approximate sites of Cas9 cleavage sites of the sgRNA used in this study are indicated by arrows. **B)** Schematic of the experimental process. C-shaped postures indicative of spontaneous C-starts / C- shaped seizures (C-SC) are indicated in two tadpoles (black asterisk). **C)** Example Sanger sequence traces with Cas9 breakpoints for both sgRNA indicated by black arrows. The most common outcomes for each guide are shown as sequence and protein schematics: rnk5 deletes 15bp leading to a loss of 5 amino acids in the bHLH DNA binding domain, indicated by the red arrow. The NLS KKRKMTK is reformed around this deletion. Rnk20 sgRNA causes a 4 bp deletion which results in a premature STOP codon (orange asterisk) and a truncated protein lacking much of the DNA binding domain is predicted.

### Ca^2+^ imaging and quantification

GCaMP6s and mCherry in CS2+ plasmid (Li *et al.* 2022) were a kind gift from Edward Ruthazer and Anne Schohl. Both were linearised with NotI HF restriction enzyme (NEB) and mRNA was synthesised with the Invitrogen mMessage mMachine SP6 kit. A total of 500 pg GCaMP6s and 250 pg mCherry mRNA was injected bilaterally into 1-cell stage *X. laevis* embryos. At stage 47, tadpoles were initially anesthetized by immersion in a petri dish containing a 1:4000 dilution of MS222 for a period of 2 minutes. Once movement ceased, tadpoles were transferred to a clean petri dish and surplus liquid removed. Tadpoles were then embedded in 40 μl 2% low melting point (LMP) agarose and immobilised by submerging in 0.1 x MMR containing 100 μM of the neuromuscular blocker pancuronium bromide (Sigma). After 5-10 minutes, fine, sharpened jewellers forceps (Dumont #5) were used to gently remove the skin above the brain, which was then covered with a thin layer of 1% LMP agarose. Prepared tadpoles were positioned beneath a fluorescence microscope (Axio Examiner D1; Carl Zeiss), and brains imaged through a 50x magnification lens (50x/0.55 DIC EC Epiplan-Neofluar, Zeiss). GCaMP6s fluorescence was excited at 480 nm by a monochromator (Polychrome V; T.I.L.L. Photonics). The excitation light intensity was reduced to 10% of its maximum. The emitted light was detected with a CCD camera (Sensicam; PCO Imaging) through a 495 nm dichroic mirror and a 505 nm long-pass filter. Pixels were binned on chip (4 x 4), resulting in an image resolution of 344 x 260 pixels. Images were acquired at a frame rate of 2 frames per second with an exposure time of 400 msec per frame. Baseline brain activity was recorded for 30 minutes for both Cas9 control and *NeuroD2* CRISPant tadpoles. Cas9 control tadpoles were subsequently exposed to 15 mM PTZ to induce acute seizures as a positive control. All tadpoles remained viable at the conclusion of the experimental procedures.

Imaging data were analysed with FiJi software (Schindelin *et al.* 2012). Ca^2+^signals were calculated as relative fluorescent change for each frame i of the recording as ΔF/F = (F_i_-F_B_)/F_B_, where F_i_ is the absolute fluorescence of the i^th^ frame and F_B_ is the background fluorescence, which was calculated as the median fluorescence of the entire recording. Using standard region-of-interest (ROI) parameters, Ca^2+^ signals from the midbrain were calculated by averaging ΔF/F % for each frame. “Large” (>5 ΔF/F %) and “small” (1-5 ΔF/F %) spikes were annotated manually. Total spike counts (small plus large) were compared between groups using a *t-*test with Mann-Whitney corrections.

### Blood-brain barrier (BBB) permeability assay

To assess BBB permeability, stage 47 tadpoles were anesthetized and embedded in LMP agarose, as for Ca^2+^ signalling, covering the brain with a thin layer. Once the agarose had set, embedded tadpoles were submerged in 0.1 x MMR and 9.2 nl of 0.1 mg/ml NaF was injected into the 4^th^ ventricle using a capillary needle (as for embryo injections). Tadpoles were protected from light following injection. A Leica fluo III dissecting microscope with GFP2 filter set was used to image the tadpole head from the dorsal side at 2, 5, 10, 20 and 30 minutes post injection using DFC7000T camera with standardized settings. Images were analysed offline using FiJi. The mean fluorescent intensity (MFI) was calculated from each image using the green channel across a defined region of interest (ROI). The ROI was located to the left side of the tadpole brain, and captures NaF dispersion across the blood-brain barrier. A 2-way repeated measure ANOVA with Tukey post hoc multiple comparisons tests of all means was used to compare the MFI, representing NaF dispersion, at 2 and 20 minutes post injection.

### Chemical treatments

#### Acute seizure induction with pentylenetetrazole (PTZ)

Healthy, normally developing stage 47 tadpoles were arbitrarily selected and placed in 0.5 ml 0.1 x MMR in separate wells of a 24- well plate. To induce acute seizures/status epilepticus as described in (Hewapathirane *et al.* 2008), the seizure-inducing drug PTZ was made up in Milli Q water (MQW) at 100 mM and 25 μl added to individual tadpole wells to give a final concentration of 5 mM. For Ca^2+^ recordings, 15 mM PTZ was used.

#### Pre-treatments with anti-seizure drug valproic acid (VPA) and the anti-inflammatory drug Losartan

Wild type tadpoles were arbitrarily divided into two groups. For each test, one group of 12 tadpoles was pre-treated for two hours with either 5 mM Sodium valproate (VPA, Sigma) or 5 mM Losartan (potassium salt, Sigma), with the other untreated group left as a control. Some of the *NeuroD2* CRISPants, selected based on spontaneous seizure behaviour, were pre-treated with 5mM Losartan prior to Ca^2+^ signal recordings and the blood-brain barrier integrity assays. Untreated *NeuroD2* CRISPants were used as controls for comparison.

### Graphs and Statistical Analysis

All graphs and statistical analyses were prepared in GraphPad Prism v10. Raw data and analysis for all figures can be found in File S1. The Shapiro-Wilk test for normality was used to confirm normal distributions, with non-parametric tests being used in the event that distributions were non-normal.

## Results

### *Xenopus laevis NeuroD2* CRISPant tadpoles exhibit spontaneous seizures with an overlapping but distinct behavioural profile compared to PTZ treated wild type tadpoles

*Xenopus* tropicalis *NeuroD2* CRISPant tadpoles were previously noted to have C-shaped contractions (Sega *et al.* 2019), similar to those elicited by exposure to epileptogenic drugs such as PTZ in wild type *X. laevis* tadpoles (Hewapathirane *et al.* 2008). *X. laevis* is allotetraploid, with around 60% of genes existing in both *L* and *S* chromosomal locations (Session *et al.* 2016). To first determine the usefulness of *X. laevis* as a genetic model for DEE, we designed sgRNA that would bind and target both copies of *NeuroD2*, to see if we could replicate the DEE75 phenotype (Figure 1A). We kept tadpoles in individual arenas so that they could not influence each other’s behaviour, and analysed video recordings to identify and quantify behaviour.

Genotyping and other assays were performed after this step (Figure 1B). Two sgRNA were designed to target the bHLH domain coding region of both *NeuroD2*.*S* and *NeuroD2*.*L*, “rnk5” resulted in a 15 bp deletion, removing 5 amino acids (aa) including part of the nuclear localisation sequence (NLS). Despite the loss of 5 aa, the consensus NLS sequence KKRKMTK was recreated around the breakpoint (Figure 1C). The second sgRNA, “rnk20”, targets the DNA binding region where the DEE75 causing variants are found in humans. This sgRNA causes frameshift InDels, most commonly a deletion of 4 bp, resulting in a truncated protein midway through the bHLH domain (Figure 1C).

Automated tracking of movement from video recordings of individual tadpoles housed in 24 well plates showed increased activity in *NeuroD2* CRISPant tadpoles (Figure 2A). Compared to wild type tadpoles induced to seizure behaviour with PTZ, which swam consistently around the periphery of the arena, *NeuroD2* CRISPant tadpoles spent more time in the centre of the well. As a control for the effect of CRISPR reagent injection on behaviour, we also tracked 12 tadpoles that had only been injected with the unloaded Cas9 protein. One of these Cas9 controls was very active, comparable to that of *NeuroD2* CRISPants, but the others appeared to behave similarly to uninjected control tadpoles, which either drifted slowly or swam for short periods around the periphery.

**Figure 2:**
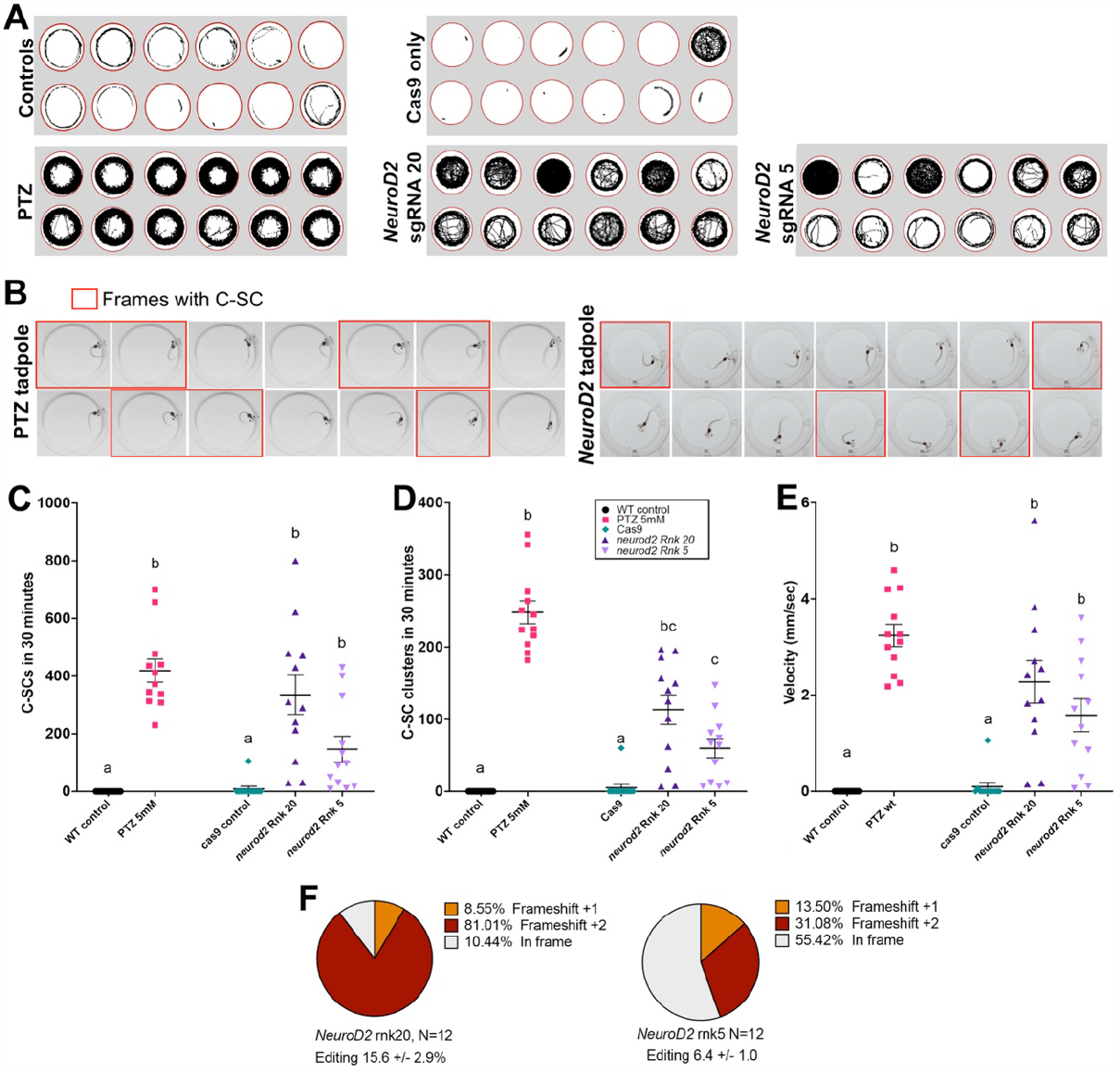
CRISPR/Cas9 edi4ng of *NeuroD2* in tadpoles results in increased activity and C-shaped contractions of the tail. **A)** Tracks showing swimming trajectories of 12 individual tadpoles in wells of a 24 well plate (indicated by red circles), recorded over 30 minutes. **B)** 14 consectutive video frames from example tadpoles, each frame is 40 msec apart. Frames where the tadpole is undergoing a C-shaped contractions (C-SC) of the tail are outlined in red. Left: wild type tadpole treated with 5mM PTZ to evoke acute C-SC; right: *NeuroD2* CRISPant tadpole with spontaneous C-SC. **C-E)** Scatter plots of behavioural data from groups of 12 tadpoles, analysed from 30 minutes of video data at 25 frames per second. Kruskal-Wallis statistical analysis with Dunns post hoc tesing of all means. Treatment groups that share the same lower case letter are not significantly different, legend in D applies to all graphs. **C)** Manually counted C-SC frames from 30 minute video recordings, **D)** Manually counted clusters of C-SC seizures **E)** Mean velocity (mm/sec). **F)** Summary of editing types detected by TIDE in the *NeuroD2* CRISPant tadpoles used to generate this data. Raw count data and statistical analyses can be found in File S1.

Previously, a type of induced seizure behaviour has been described in *X. laevis* tadpoles as “C-shaped seizure” where the trunk curls to one side in an extreme posture often resulting in the head touching the tail tip (Hewapathirane *et al.* 2008). Frame-by-frame analysis of video capture shows that this behaviour can be elicited by adding 5 mM PTZ to the tadpole’s swimming water (Figure 2B). While C-shaped postures were frequently seen in both PTZ-induced and *NeuroD2* CRISPant tadpoles, PTZ tadpoles were more likely to hold the posture for consecutive frames, indicating an acute convulsive state that resembles status epilepticus (Figure 2B). PTZ-elicited seizures have been confirmed by both electrophysiological field potential recordings from the tadpole brain and Ca^2+^ signalling (Hewapathirane *et al.* 2008). Here, we refer to “C-shaped contractions” (C-SC) to describe the behaviour seen in *NeuroD2* CRISPant tadpoles (Video S1). Both individual frames with C-SC and C-SC clusters (where multiple C-SCs were detected, with an interval of less than 2 seconds between events), were manually counted from the same tadpoles for the 30 minutes of video recorded footage (Figure 2C, 2D). PTZ induced tadpoles had the most C-SC (419 +/- 40.2), compared to 335 +/- 68.1 for rnk20 and 146 +/- 44.6 for rnk5 *NeuroD2* CRISPant tadpoles (Figure 2C, Video S2). The truncating rnk20 *NeuroD2* CRISPant tadpoles therefore had an average of 2.3 times as many C-SC as the 5 aa deletion-causing rnk5. Kruskal-Wallis analysis of the C-SC data, with Dunn’s post hoc testing of means, showed that rnk5 mean C-SC count was not increased over that of controls, while for rnk20, the higher mean C-SC count was not significantly different from PTZ-induced tadpoles. A similar relationship was seen with C-SC clusters (Figure 2D), except that rnk5 was shown to have significantly more C-SC clusters than controls. Both sgRNA had significantly lower numbers of C-SC clusters compared with PTZ-treated tadpoles.

We also analysed the same videos with TopScan and found that mean velocity (mm/sec) was an excellent indicator of seizure activity, correlating strongly with C-SC counts (Figure 2E, Figure S2B). Conversely, TopScan automated scoring of C-SC was found to be inaccurate for *NeuroD2* CRISPant tadpoles ( Figure S2A) On average, the truncating rnk20 *NeuroD2* CRISPant tadpoles swam 25 times faster than cas9 injected controls, and the 5aa deletion rnk5 *NeuroD2* CRISPant tadpoles swam 18 times faster than the controls. PTZ kindled tadpoles swam fastest with a mean velocity of 3.24 +/- 0.23 mm/sec, compared to 2.27 +/- 0.44 for Rnk20 and 1.58 +/- 0.34 for rnk5 *NeuroD2* CRISPant tadpoles. Editing for both *NeuroD2* CRISPant groups was confirmed from Sanger sequence trace analysis using TIDE, with all tadpoles showing editing. For rnk20 tadpole editing varied from 3.5% to 33.3% with a mean of 15.6 +/- 2.9%, and for rnk5 the range was 2.8-12.3, mean 6.4 +/- 1.0 (Figure 2F). Because of the more obvious effect of the rnk20 edits on the NeuroD2 protein, as well as higher average editing, we proceeded with this sgRNA for further investigations.

### *NeuroD2* CRISPant tadpoles show strong, prolongued, and widepread Ca^2+^ signalling activity in the brain

To determine if the C-SC and faster swimming behaviours we detected in CRISPants were indicative of abberant brain signalling, we used the genetically encoded Ca^2+^ indicator fusion protein GCaMP6s to record Ca^2+^ signals from tadpole brains at stage 47 (Figure 3). Single cell embryos were injected bilaterally with mCherry and GCaMP6s mRNAs in addition to the CRISPR reagents (Figure 3A). Expression of mCherry in both halves of the neural plate at stage 16 was used to confirm delivery of reagents. At stage 47, tadpoles were first video recorded, and individuals with prominent C-SC were selected and mounted for live monitoring of Ca^2+^ signals over 30 minutes. Background recordings, or those from control, Cas9 only injected tadpoles (both N=12), showed a generally low level of Ca^2+^ signals (observed as GFP fluorescence), but spikes of activity could still be detected (Video S3). Some of these control tadpoles were then exposed to 15 mM PTZ and recording continued for a further 30 minutes. PTZ significantly increased Ca^2+^ signal intensity (median cross entire tadpole 6.7 +/- 1.1%) compared to 0.7 +/- 0.1% pre-PTZ exposure, or to 2.2 +/- 0.3% in untreated siblings at the same timepoint (Figure 3D, Figure S3, Video S4). The average total number of large and small spikes was also significantly higher in PTZ treated tadpoles than sibling controls (7.29 +/- 1.78, Figure 3E). *NeuroD2* sgRNA rnk20 CRISPants, on the other hand, showed additional strong, widespread signals that originated from a focal point before spreading across the entire brain on both hemispheres (Video S5). These episodes, which generated large Ca^2+^ spikes, lasted between three to five minutes in some cases and may indicate epileptogenic activity (Figure 3C, Figure S3). Even inbetween these episodes, Ca^2+^ signalling in CRISPant tadpoles was elevated, with increased numbers of smaller Ca^2+^ spikes. Total Ca^2+^ spike counts over 30 minutes was significantly higher in the *NeuroD2* CRISPant group (mean 23.42 +/- 4.2, N=12) than in Cas9 injected controls (9.67 +/- 2.07, N=12) (Figure 3F). All CRISPant tadpoles were confirmed with editing in the range 7.7 to 42.9%, and 100% of edits resulted in frameshifts (Figure 3G).

**Figure 3:**
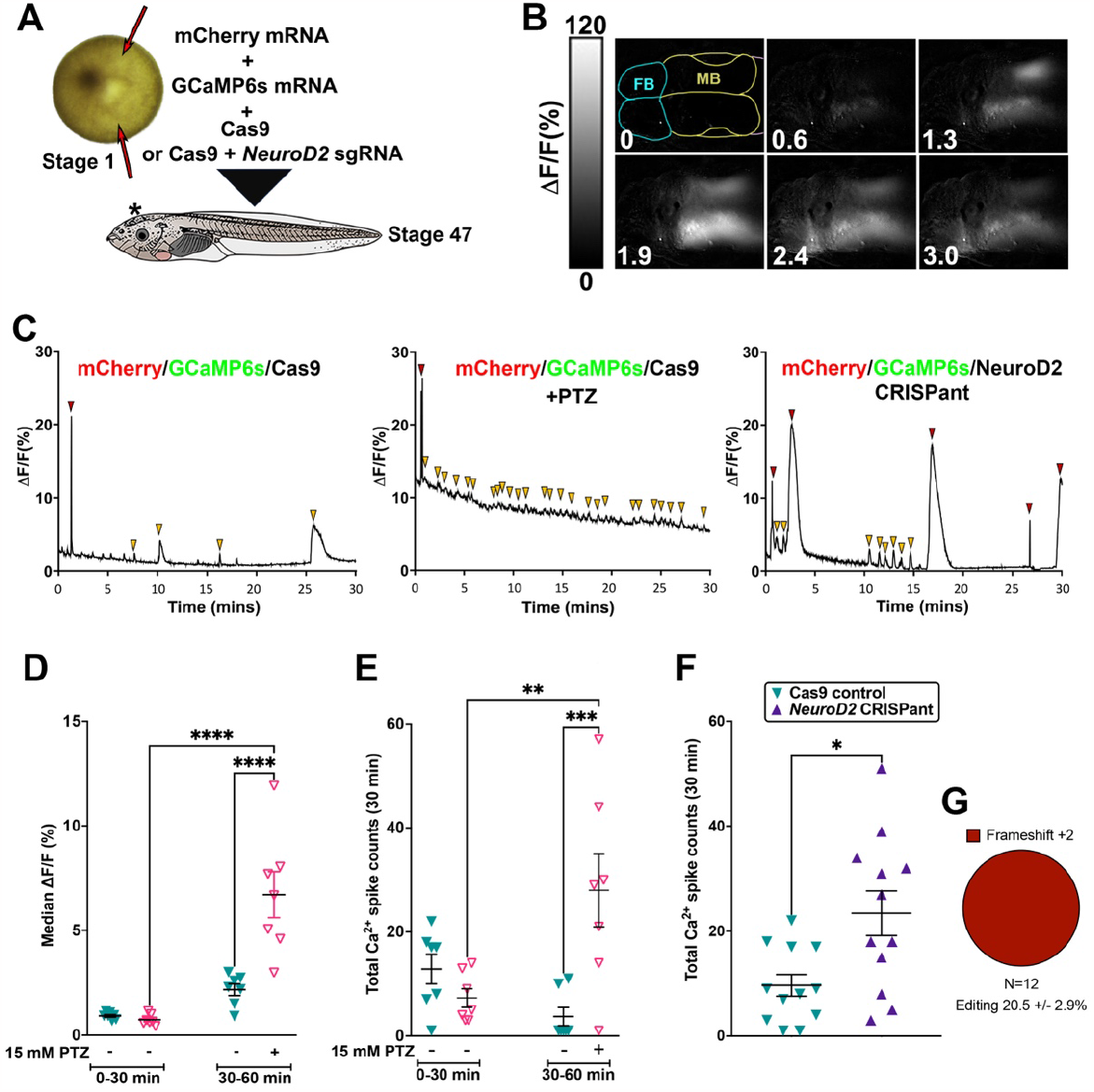
*NeuroD2* CRISPant tadpoles show strong, prolongued, and widepread Ca^2+^ signals in the brain. **A)** Experimental design. Fertilised *X. laevis* embryos at the one cell stage were injected twice (red arrows) on either side of the female nucleus (light spot), avoiding the ventrally located sperm entry point (dark spot). mCherry mRNA was used to confirm injection, GCaMP6s mRNA encodes a Ca^2+^-sensing fusion protein that can detect action potentials, emitting strong fluorescence in the GFP channel signals (Li *et al.* 2022). Controls were only injected with Cas9 protein, and CRISPant embryos were injected with Cas9 with *NeuroD2* sgRNA rnk20 loaded. Embryos were raised to stage 47. **B)** Example still frames taken every 0.64 minutes from a live *NeuroD2* CRISPant tadpole brain (TILLvisION 4.0). CaMP6s fluorescence (lighter colour = higher intensity) can be seen across large regions of the brain. Scale bar indicates Ca^2+^ signal intensity (%). Forebrain (FB) hemispheres outlined in blue, midbrain (MB) in yellow, hindbrain in pink is mostly out of shot. Time in minutes is indicated in the bottom left. **(C)** Examples of Ca^2+^ signals in the midbrain optic tectum (mean fluorescent intensity across 30 minutes), in control, PTZ induced wild type tadpoles and *NeuroD2* CRISPants. Red arrowheads indicate large spikes and gold arrowheads small spikes. **(D-F)** Scatterplots showing individual tadpoles as triangles, black horizontal bars are means and error bars show SEM. **D)** Median midbrain Ca^2+^signals in two groups of N=7 Cas9 control tadpoles. One group was exposed to 15 mM PTZ to induce seizure activity in the second recording time window (30-60 minutes). 2-way ANOVA, post-hoc test of all means (uncorrected Fisher’s LSD) **** p<0.0001 **E)** Comparison of total (large and small) Ca^2+^ spikes in the same tadpoles and groups as in panel D, 2-way ANOVA, ** p<0.01, *** p<0.001. **F)** Comparison of total Ca^2+^ spikes in Cas9 control tadpoles vs. *NeuroD2* CRISPants (N=12) Mann-Whitney Test * p<0.05). **G)** Summary of editing types detected by TIDE in the *NeuroD2* CRISPant tadpoles used to generate this dataTrace data for all tadpoles in the study can be found in Figure S3, raw data and statistics in File S1.

### *NeuroD2* CRISPant tadpole brains appear morphologically normal, but have a leaky blood- brain barrier (BBB)

We next looked to see if *NeuroD2* CRISPant tadpoles would develop gross brain morphological defects. We first utilised the unilateral mutant method pioneered by Willsey et al in *X. tropicalis*, in a study of autism associated genes (Willsey *et al.* 2021). CRISPR reagents (Cas9 loaded with *NeuroD2* rnk20 sgRNA) were injected into one blastomere at the two cell stage of development, along with mRNA for nGFP. This results in one side of the embryo inheriting both the CRISPR reagents and nGFP, while the other side remains wildtype. At stage 19, the resulting embryos were sorted into left and right unilateral CRISPants, based on the GFP location. Embryos were raised to stage 47 and tadpoles photographed in order to compare the morphology of the left and right brain hemispheres. No differences in gross morphology could be detected in either 16 left or 19 right-sided unilateral tadpoles, obtained from two sibships (Figure S4).

We also measured the permeability of the blood-brain barrier (BBB) in both *NeuroD2* CRISPants and PTZ acute seizure induced tadpoles. NaF dye has been previously shown to be prevented from exciting the brain at stage 55 in uninjured tadpoles, indicating an intact BBB (De jesus andino *et al.* 2016). Adapting the method used in a recent study using *X. laevis* tadpoles to model traumatic brain injury (Spruiell eldridge *et al.* 2022) for our younger tadpoles, we injected 9.2 nl of 0.1 mg/ml sodium fluorescein (NaF) into the 4th ventricle (Figure 4A) at stage 47. BBB permability was then measured by tracking dye distribution over 30 minutes (Figure 4B). PTZ treatment to elicit status epilepticus did not result in increased NaF loss from the brain compared to WT or Cas9 only injected controls at either timepoint. In contrast, *NeuroD2* CRISPants showed significantly higher NaF fluorescence outside the brain loaded with *NeuroD2* rnk20 sgRNA) were injected into one blastomere at the two cell stage of development, along with mRNA for nGFP. This results in one side of the embryo inheriting both the CRISPR reagents and nGFP, while the other side remains wildtype. At stage 19, the resulting embryos were sorted into left and right unilateral CRISPants, based on the GFP location. Embryos were raised to stage 47 and tadpoles photographed in order to compare the morphology of the left and right brain hemispheres. No differences in gross morphology could be detected in either 16 left or 19 right-sided unilateral tadpoles, obtained from two sibships (Figure S4).

**Figure 4:**
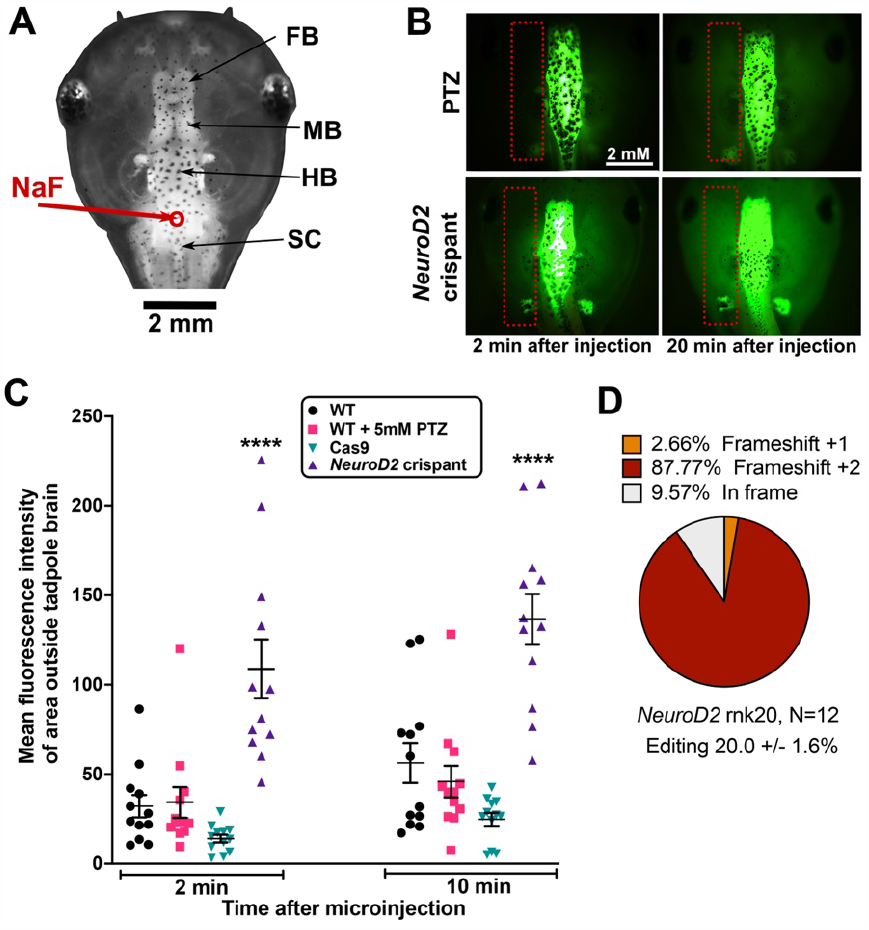
*NeuroD2* CRISPant tadpoles have a comparativly leaky blood-brain barrier. **A)** Dorsal view of stage 47 tadpole head to show the parts of the brain and site of Na fluorescien (NaF) injection into the hindbrain ventricle (red circle and arrow). FB: forebrain, MB: midbrain, HB:hindbrain, SC, spinal cord. **B)** Examples of NaF dye injected tadpoles at 2 and 20 minutes after injection, visualised with GFP2 channel. Orientation of the tadpole head as in (A). Red dotted rectangles show the area outside the brain that was used to calculate mean fluorescent intensity. Scale bar in top left applies to all panels. Top, wild type (WT) tadpole treated with 5 mM of seizure-inducing drug PTZ for 2 hours prior to dye injection. Bottom, untreated *NeuroD2* CRISPant, sgRNA rnk20. **C)** Scatter plot showing MFI (dye leakage outside the brain) at 2 and 20 minutes post NaF dye injection for N=12 tadpoles per group, 2-way ANOVA with Tukey post-hoc analysis, **** p<0.0001. Cas9 indicates tadpoles injected with Cas9 protein, but no sgRNA. **D)** Summary of tadpole editing in the *NeuroD2* CRISPant group, as confirmed by Sanger sequencing and TIDE analysis. Raw data can be found in File S1 and Figure S5.

We also measured the permeability of the blood-brain barrier (BBB) in both *NeuroD2* CRISPants and PTZ acute seizure induced tadpoles. NaF dye has been previously shown to be prevented from exciting the brain at stage 55 in uninjured tadpoles, indicating an intact BBB (De jesus andino *et al.* 2016). Adapting the method used in a recent study using *X. laevis* tadpoles to model traumatic brain injury (Spruiell eldridge *et al.* 2022) for our younger tadpoles, we injected 9.2 nl of 0.1 mg/ml sodium fluorescein (NaF) into the 4th ventricle (Figure 4A) at stage 47. BBB permability was then measured by tracking dye distribution over 30 minutes (Figure 4B). PTZ treatment to elicit status epilepticus did not result in increased NaF loss from the brain compared to WT or Cas9 only injected controls at either timepoint. In contrast, *NeuroD2* CRISPants showed significantly higher NaF fluorescence outside the brain at both the 2 minutes (7.7 x higher) and 20 minutes timepoints (5.5 x higher), compared to Cas 9 controls (Figure 4C). All CRISPant tadpoles were subsequently confirmed as edited (Figure 4D). This rapid loss of NaF from brain of *NeuroD2* CRISPants indicates that the BBB is inherently leaky in these tadpoles.

### Pretreating tadpoles with the anti-inflammatory drug losartan offers short term protection from chemically induced or genetic seizure activity

Anti-seizure drugs (ASD) are front-line medications used in epilepsy to prevent seizures from occurring. However, there is a current lack of effective seizure controlling medication for treatment of DEE. Drug re-purposing for epilepsy, particularly the use of anti-inflammatory and anti-oxidant medications with known, and manageable, side effect profiles, is increasingly desirable (for recent reviews, see (Radu *et al.* 2017; Klein *et al.* 2020; Loscher 2020; Pawlik *et al.* 2021). We tested two drugs, the commonly used anti-seizure drug VPA, and the anti- inflammatory drug Losartan, to determine the potential for seizure reduction. Losartan is a commonly prescribed drug used to lower blood pressure. It acts by inhibiting angiotensin II type I receptor antagonist, leading to up-regulation of the protease thrombospondin1 (TSP1). TSP1 can activate the pro-protein form of the secreted paracrine factor TGFβ (Bar-klein *et al.* 2014). Losartan has been previously reported to reduce the development of chronic seizures in rodent models of traumatic brain injury (Bar-klein *et al.* 2014; Tchekalarova *et al.* 2016; Hong *et al.* 2019), but to our knowledge this drug has not been tested in models of genetic epilepsy such as DEE75.

Tadpoles were pre-treated for 30 minutes with 5 mM VPA, 5 mM Losartan or no drug, before adding 5 mM PTZ to the swimming water to induce seizures. The number of C-SC was counted in two 10-minute windows, 20-30 minutes and 50-60 minutes after adding PTZ (Figure 5A, B). At the earlier timepoint, the VPA pre-treatment tadpole cohort had significantly fewer mean C-SCs than controls (Figure 5A, control mean C-SC: 103.2 +/- 11.4, VPA mean C-SC 17.3 +/- 3.8). The losartan pre-treatment cohort also had significantly fewer C-SCs than controls, (Figure 5B, control mean C-SC 30.8 +/- 7.6, Losartan mean C-SC 8.5 +/- 3.2). Despite their different models of action, both drugs were able to offer some protection from PTZ induced seizure behaviour, but the protection afforded by losartan did not extend to the later timepoint, suggesting it is short-lived in the acute seizure model. We also found that tadpoles in the VPA pre-treatment group were significantly slower swimmers, whereas pre-treatment with losartan had no effect on mean swimming velocity (Supplementary Figure S7).

**Figure 5.**
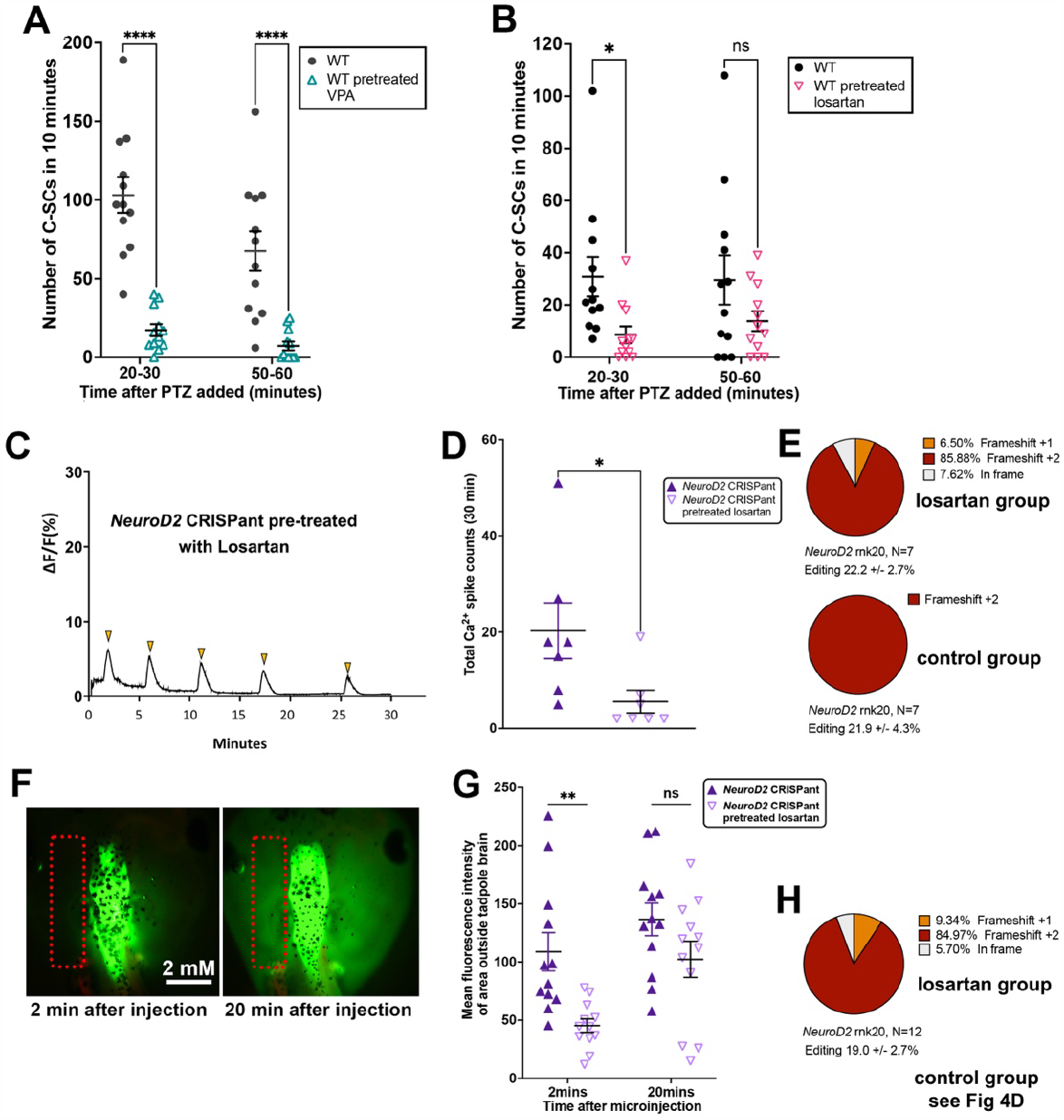
Pre-trea4ng tadpoles with the an4-inflammatory drug losartan offers short-term protec4on from chemically induced or gene4c seizure ac4vity. **A,B)** Scatter plots of C-SC events in stage 47 wild type tadpoles recorded in 10 minutes either 20 minutes after induction of seizures with 5 mM PTZ or 50 minutes after. Analysis by 2-way ANOVA with Sidak’s multiple comparisons test of means, * p<0.5, **** p<0.0001, ns non significant. **A)** Effect of pre-treatment of tadpoles with 5 mM of the anti-seizure drug VPA, compared to no pretreatment, N=12 both groups. **B)** Effect of pre-treatment of tadpoles with 5 mM of the anti-inflammatory drug losartan, compared to no pretreatment, N=12 both groups. **C)** Example Ca ^2+^ signal trace signalling in the midbrain optic tectum of a *NeuroD2* CRISPant tadpole pre-treated with 5mM losartan, with spikes of Ca ^2+^ activity indicated by gold arrowheads. **D)** Scatterplot of total Ca ^2+^ spikes, detected over 30 minutes, for two groups of N=7 *NeuroD2* CRISPant tadpoles with or without losartan pre-treatment. Mann-Whitney test, * p<0.05. **E)** Summary of editing for each group of n=7 tadpoles in panel D. All CRISPants were edited, and levels were not different between the groups (p=0.96, unpaired t-test). **F)** *NeuroD2* CRISPant tadpole heads showing distribution of sodium fluorescein (NaF) dye 2 and 20 minutes after injectio in to the 4th ventricle. Red dotted rectangle indicates the area outside the brain used to measure escaped NaF. **G)** Scatterplot comparing mean fluorescence intensity in the brain at 2 and 20 minutes post NaF injection, for two groups of 12 *NeuroD2* CRISPant tadpoles, where one group was pretreated with 5 mM Losartan. Analysis by 2-way ANOVA with Sidak’s multiple comparisons test of means, ** p<0.01, ns=non significant. **H)** Pie chart showing editing breakdown the 12 tadpoles the losartan group in panel G. Controls are shown in Fig. 4D All CRISPants were edited, and levels were not different between the groups (p=0.76, unpaired t-test). Raw data can be found in File S1 and Figure S5.

We then investigated whether Losartan pre-treatment could be used to reduce the aberrant brain signalling activity seen in our *NeuroD2* CRISPants. We generated CRISPant tadpoles with mRNA encoded GCaMP6s and arbitrarily assigned them to a Losartan pre-treatment or control groups of N=7, to observe the effect GCaMP6s fluorescence (indicating Ca^2+^ signalling) in the stage 47 brain (Figure 5C, D). Significantly fewer Ca^2+^ spikes were seen in the group of *NeuroD2* CRISPant tadpoles treated with 5 mM Losartan for 2 hours prior to recording, nearly a 4-fold reduction. Untreated CRISPant tadpoles had a mean of 20.3 +/- 5.8 spikes in 30 minutes compared to the Losartan pre-treated batch with 5.6 +/-2.4 spikes in 30 minutes. Editing was confirmed for all tadpoles, with no difference between treatment groups (N=7 per group, Figure 5E). We also examined the effect of 2 hours of pre-treatment with 5 mM Losartan on the BBB integrity of *NeuroD2* CRISPants using the NaF assay, comparing to controls from Figure 4C. *NeuroD2* CRISPant tadpoles in the Losartan pre-treatment group were initially better able to retain the NaF dye in the brain (MFI outside the brain was 108.80 +/- 16.43 for controls and 45.32 +/- 5.76 for the Losartan group). However, analysis at 20 minutes after injection showed there was no difference between the two groups (mean 136.6 +/- 13.98 for controls and 102.3 +/- 15.53 for Losartan). Editing was again confirmed for all tadpoles, with no difference between treatment groups (N=12 per group, Figure 5H and 4D).

### Gene editing efficiency correlates positively to the number of C-shaped contractions or clusters of contractions observed in *NeuroD2* CRISPants

Patients with DEE-75 resulting from de novo, non-synonymous changes to the *NeuroD2* coding region would still be expected to have one functional copy of the *NeuroD2* gene. Therefore, the amount of functional *NeuroD2* protein during brain development would be expected to be around 50% of that of unaffected individuals. The amount of editing seen in tadpoles varied from 3 to 43%, so we tested whether there is a correlation between the efficiency of gene editing and the severity of the seizure-like phenotype. We used two measures for severity: seizure frequency (C-SC per minute) and seizure cluster frequency (C-SC clusters per minute, with a cluster being defined as one or more C-SCs separated by at least two seconds from the previous or next C-SC). Seizure frequency was positively correlated to editing efficiency (Figure 6A; linear regression, log-transformed editing efficiency, adjusted R^2^= 0.142, p = 0.04). Seizure cluster frequency was also positively correlated to editing efficiency, but the correlation was twice as strong (Figure 6B; linear regression, log-transformed editing efficiency, adjusted R^2^= 0.29, p = 0.004). This stronger correlation indicates that seizure clusters are a more sensitive measure for seizure severity than seizure frequency. For both severity measures (seizure frequency and seizure cluster frequency) the type of sgRNA (rnk5 or rnk20) did not affect the correlation between editing efficiency and seizure severity (linear model with sgRNA rnk [rnk5 and rnk20] as explanatory variable, p > 0.2).

**Figure 6:**
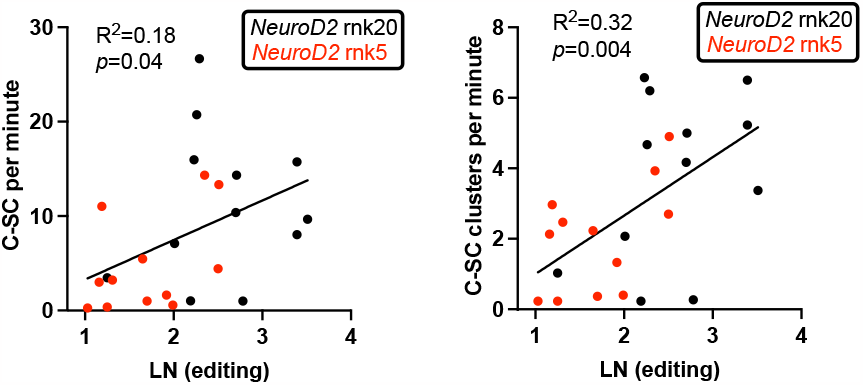
Clusters of C-SC correlate more strongly to *NeuroD2* gene edi4ng levels than counts of individual C-SC. Regression plots of C-SC (left) or clusters of C-SC (right) per minute against log transformed editing % (LN(editing %)) of *NeuroD2*. Each dot represents an individual tadpole (N=24) with *NeuroD2* sgRNA rnk5 indicated in red and sgRNA rnk20 in black. R^2^ indicates correlation and *p-*value indicates significance. Raw data can be found in File S1.

## Discussion

### Tadpole behaviour analysis can be useful phenotyping tool for pre-clinical models of DEE

Tadpoles of *X. laevis* frogs develop along a well-defined series of morphologically described stages (Nieuwkoop And faber 1967). Since developmental rate is dependent on temperature for this species, we chose to evaluate behaviour at stage 47. If tadpoles are not fed, they will remain at this stage, whereas earlier stages are more transient. We observed a considerable amount of batch variability in baseline behaviour in this study, with Cas9 or uninjected control tadpoles barely moving at all (Figure 1) compared to velocities of 4.8 mm/sec in Figure 4. Currie et al looked at the development of swimming behaviour in *X. laevis* tadpoles (Currie *et al.* 2016). They measured swimming velocity in a 50 mm arena and showed that tadpoles began swimming at stage 45, but by stage 47 they were active almost continuously and swam at a mean velocity of 10 mm/sec. While we used a smaller arena size (17 mm diameter) this is unlikely to account for the magnitude of these differences. It is possible that our practice of withholding food contributes, and further work will be needed to ascertain if this is the case. Alternatively, our tadpoles could be swimming less due to exposure to strong light, which is used for video recording. *X. laevis* tadpoles sense light via the pineal gland, and swim more in darkness (Jamieson And roberts 2000).

Automation of behavioural characteristics using TopScan in 24 well plate arrayed tadpoles has been previously described (Panthi *et al.* 2023). In *NeuroD2* CRISPants, mean swimming velocity was elevated compared to controls, and is a good indicator of hyperactivity, also described in mouse *NeuroD2* knockouts (Runge *et al.* 2021). Swimming track data highlighted differences in the occupancy of the arena, with *NeuroD2* CRISPants tending to cross the centre more frequently than PTZ-induced acute seizure tadpoles. This also seems to mirror the mouse knockout (Runge *et al.* 2021). While others have reported a recapitulation of the PTZ-induced behavioural phenotype in *NeuroD2* CRISPants of the diploid *X. tropicalis*, in isolated *X. laevis* tadpoles we were able to detect differences in the two behaviours, with CRISPants more often just showing C-SC in single frames and having more periods of inactivity between C-SC clusters. For PTZ induced tadpoles, TopScan was able to identify C-SC accurately, using elongation ratio thresholds (altered length/width), as confirmed by manual counts. On the other hand, spontaneous C-SC of the CRISPants were not reliably detected, and needed to be manually scored on a frame-by-frame basis, possibly due to the more transient nature of the C-shaped postures. We note this difference from Sega et al’s model of DEE75 in *X. tropicalis*, where much stronger seizure activity was reported with tadpoles spending up to half of their time in C-SC at stage 47 (Sega *et al.* 2019), when multiple tadpoles were in the same dish. This difference could be due to our tadpoles being individually housed, with no opportunity to influence each other’s behaviour. We did however, see a strong, significant correlation between increased mean swimming velocity and manually counted C-SC, indicating that seizure behaviour can be assessed in this model by automatically measuring this parameter, as previously shown for induced seizures (Panthi *et al.* 2023).

### Both truncating edits and the deletion of amino acids in the DNA binding domain result in seizure-like behaviour

We used two sgRNA with different predicted outcomes in this study. For sgRNA rnk20, we obtained sequencing data for 20 stage 14 embryos and 55 stage 47 tadpoles after phenotyping (Figure S1A). InDelphi predicts a predominant 4bp deletion, which was confirmed from TIDE analysis of Sanger sequence traces in all 55 sequenced tadpoles, as well as in 18 of 20 embryos. The second highest prediction by InDelphi for this target is a 13 bp deletion, which was detected in just 4/20 embryos (20%) and 14/55 tadpoles (25%). We conclude that the most common outcome for this sgRNA is a deletion of 4 bp which results in a change of amino acid residue at position 135, followed by a stop codon (nonsense mutation). The outcome of the less common 13 bp deletion is the same, as both deletions result in +2 frameshift. Any protein product formed from rnk20 sgRNA editing is therefore likely to be truncated just 3’ of the NLS, with a shortened bHLH DNA binding domain (Figure 1C). We expect this to represent a mosaic loss of function phenotype. In mice, loss of *NeuroD2* results in ataxia and death by postnatal day 14 (Olson *et al.* 2001). For the rnk5 sgRNA, InDelphi predicts a 15 bp deletion, which was observed in half of our tadpoles (5/10, Supplementary Figure S1B). This creates a deletion of 5 amino acids in the region of the NLS, but the NLS itself is preserved (Figure 1C). The outcome of this *NeuroD2*Δ5 deletion is less clear. Perhaps surprisingly, this sgRNA was also able to induce C-SC behaviour, even with relatively low editing levels. Potentially, the loss of 5 amino acids in the bHLH domain may affect the structure of the transcription factor, resulting in reduced function. Our results suggest that embryos are very sensitive to *NeuroD2* protein levels during early development. Sega et al found that overexpression of wild type *NeuroD2* led to the production of ectopic primary neurons, whereas the DEE-75 variant replicating *NeuroD2* alleles were either less active or not active (Sega *et al.* 2019). This suggests that DEE-75 is caused by loss of function of *NeuroD2*, and patients are haploinsufficient. Haploinsufficiency is also supported by the mouse model (Runge *et al.* 2021). Here, we show that even a relatively small decrease in functional *NeuroD2* results in a detectable C-SC phenotype. The two original DEE-75 variants, Glu130Gln and Met134Tyr, are predicted to alter DNA binding (Sega *et al.* 2019), and both have been linked to DEE in at least one other patient (Runge *et al.* 2021; Demarest *et al.* 2022; Sakpichaisakul *et al.* 2022). However a third variant in the DNA binding domain, Arg129Trp, was identified in a patient with neurodevelopmental delay but no epilepsy (Runge *et al.* 2021). Two further variants, Leu163Pro (inside bHLH domain) and His268Gln have also been associated with neurodevelopmental phenotypes/autism, but not with seizures (Mis *et al.* 2021; Runge *et al.* 2021).

### C-shaped contractions resemble the Mauthner neuron-mediated C-start responses of larval fish and amphibians

Aquatic vertebrates such as fish and larval frogs (tadpoles) have a unique pair of giant axon neurons called Mauthner neurons. In zebrafish, one Mauthner neuron on each side of the animal projects its axon from the 4th rhombomere of the hindbrain to the spinal motor neurons on the contralateral side (Kimmel *et al.* 1974). The giant axons make the Mauthner neurons capable of rapid responses. Typically, the input to the Mauthner neurons is either from the nearby sensory auditory afferent neurons or from the vibration sensing lateral line cells (Sillar 2009). In *Xenopus* tadpoles, Mauthner neurons have been associated with a stereotypical fast escape response termed the C-start (Sillar And robertson 2009). In teleost fish, Mauthner neurons are well established as the command neurons for the C-start escape response (Sillar 2009). The C-shaped seizures described by (Hewapathirane *et al.* 2008; Sillar And robertson 2009; Sega *et al.* 2019) may reflect inappropriate firing of the Mauthner neurons, resulting in a posture that mimics that of the C-start. In support of this, Zarei et al found that removal of one otic vesicle (ear) resulted in all the C-start bends going in the same direction instead of 50:50 left or right (Zarei *et al.* 2017) We found that our C-shaped contractions were of longer duration than the C-starts induced by temperature stress (Sillar And robertson 2009), in stage 42 tadpoles, which last about 40 msec in total, or stimulated C- starts in stage 46 tadpoles, which lasted around 30 msec (Zarei *et al.* 2017). PTZ-induced C-SC were seen to last for about 160 msec, and the spontaneous C-SC of *NeuroD2* CRISPants 40-80 msec. In future, it may be possible to target GCaMP to Mauthner cells to see if these neurons are indeed activated in C-SC.

### The tadpole model reveals that disrupted brain activity and a leakier blood brain barrier results from haploinsufficiency of *NeuroD2*

Tadpole models of induced acute seizures had been previously described (Hewapathirane *et al.* 2008; Bell *et al.* 2011). The use of tadpole CRISPants to confirm a novel DEE resulting from de novo human variants that result in loss of function of one copy of *NeuroD2* (Sega *et al.* 2019) additionally demonstrated the utility of this model in functional studies of genetic epilepsy. Here, we have demonstrated that the allotetraploid *X. laevis*, which is better established as a neuroscience model, can also be used for genetic epilepsy. Ta et al., 2022 recently provided a comprehensive transcriptome of brain development in *X. laevis*, interrogating this data shows that only the *NeuroD2*.S homeologue is expressed in the brain. Expression is mostly in the forebrain and midbrain, and midbrain expression was seen to increase steadily from stage 44 to stage 61 (Ta *et al.* 2022).

Sega et al showed that C-SC, resembling those seen in PTZ induced epilepsy, can be observed from stage 43, increasing in frequency until stage 47. Here, we have focussed on stage 47 tadpoles as this provides a natural stalling point in tadpole development in the absence of feeding, allowing time for multiple analyses. Knockout of *NeuroD2* in a mouse model was previously shown to result in microcephaly, ataxia, and death at approximately 2 weeks (Olson *et al.* 2001). In a more recent examination of *NeuroD2* knockout mice focussing on the cerebral cortex, Runge et al showed altered corticogenesis, excitatory synapse density and turnover, resulting in hyperexcitable layer 5 neurons (Runge *et al.* 2021). Social interactions, stereotypical behaviours, hyperactivity and occasional seizures were noted in both homo- and heterozygote KO mice. Both behavioural aspects and seizures were recapitulated in conditional knockouts of *NeuroD2* in forebrain excitatory neurons, but *NeuroD2* has a much broader expression in both the human brain (Human Protein Atlas) and the tadpole brain (Ta *et al.* 2022). We observed seizures in all edited tadpoles, which may be more obvious due to continuous monitoring of the isolated animals. Ca^2+^ signalling observations showed widespread neuronal hyperactivity, confirming that C-SC and hyperactivity are accompanied by an underlying aberrant neuronal signalling. *NeuroD2* was also found to be significantly decreased at both transcript and protein level in a model of Zika virus induced microcephaly (Fujimura *et al.* 2023). We did not observe any effect on gross brain morphology in the tadpole model, since unilateral CRISPants had symmetrical brains. The integrity of the blood-brain barrier is linked to the development of seizures in acquired epilepsy (reviewed in (Gorter *et al.* 2015), but has not to our knowledge been linked to genetic DEE. Our work suggests that the BBB may be less effective at stage 47. The BBB has not been investigated in NeuroD2 pathology before, but in one report of a child with *NeuroD2* M134T variant, and one with E130Q, the corpus callosum was noted to be thin in MRI reports (Sega *et al.* 2019; Demarest *et al.* 2022). Interestingly, PTZ treatment over the same timeframe had no effect on BBB integrity, so this aspect is most likely neurodevelopmental rather than result of seizure activity in the brain. Losartan has been shown to attenuate BBB permeability in status epilepticus induced by pilocarpine (Hong *et al.* 2019).

In the initial report of DEE75, neither children’s seizures could be controlled by treatment with ACTH, prednisolone or vigabatrin and a number of other ASD (Sega *et al.* 2019). Seizure freedom was reported for the patient with the more severe E130Q variant by using the ketogenic diet, and for the M134T patient by means of vagal nerve stimulation (Sega *et al.* 2019). This highlights the need to explore alternative mechanisms of treating DEE. Angiotensin II receptor transcripts were found to be significantly elevated in a rat model of epilepsy (Pereira *et al.* 2010). These authors first demonstrated the anti-convulsive effect of losartan, an angiotensin II receptor antagonist anti-hypertensive drug, in their model. Losartan may protect from epileptogenesis by preventing TGF-β signalling in astrocytes (BAR- KLEIN *et al.* 2014). Here we have shown that pre-treatment of CRISPant tadpoles with the losartan both improves the integrity of the BBB and reduces the number of Ca^2+^ spikes in our tadpole model. Losartan is one of several non ASD that have been shown to have potential in the treatment of acquired epilepsy (reviewed in (Du *et al.* 2016). A recent retrospective study showed significantly reduced epilepsy incidence in a cohort of arterial hypertension patients from Germany who were being treated with six different angiotensin II receptor blockers (Losartan, Valsartan, Telmisartan, Olmesartan, Candesartan, Irbesartan). The same study showed that Losartan alone associated with a significantly lower incidence of epilepsy (Doege *et al.* 2022). Losartan was also effective at reducing seizures and associated neuronal damage in a mouse model of hypertension and epilepsy (Tchekalarova *et al.* 2016). Conversely, (REYES-GARCIA *et al.* 2019) resported that Losartan did not reduce epileptiform activity in resected brain slices from human patients of drug resistant epilepsy. Since the effect on the BBB integrity in the tadpole model was eventiually overcome, Losartan pharmacokinetics may play a role in this variability.

### Conclusions and Limitations of the study

We have increased the toolkit for assessing seizure activity in *Xenopus* tadpole models of DEE, by using automated detection of mean tadpole velocity and live imaging of hyperactive brain activity using Ca^2+^ imaging. We have also identified a possible failure of BBB integrity or development in DEE75, and have identified Losartan as being a potentially useful drug for reducing seizures. As DEE is genetically diverse, it remains to be seen whether BBB integrity is a common associated feature, or whether Losartan may be generally useful in preventing epileptogenesis in DEE. Even though our colony has been closed for nearly 20 years, tadpoles come from a genetically diverse population of *X. laevis* and not from an isogenic line. Taken alongside CRISPR/Cas9 variation in editing and the likelyhood of mosaicism, this means that even with sibling controls, we do note a considerable variation in tadpole outcomes. Since CRISPants are mosaic, our editing information for tadpoles may not perfectly reflect the situation in the brain. In future, it may be more informative to assess editing in extracted brains, rather than whole tadpoles. Scoring of C-SCs had to be done manually and could not be reliably automated with TopScan. A mechanism for detecting and tallying these rapid C- start like movements would be advantageous in developing these models for use in pre- clinical drug screening.

## Data Availability

All data presented are in the manuscript or in the supplementary data (Figures S1-S7 and Data File S1). Videos S1-S5 are linked to FigShare.

## Contribution roles

Conceptualization: CB, PS, SB; Data curation CB, SB; Formal analysis and visualization: SB, CB, PS; Investigation SB; Funding acquisition, Project administration, Supervision: CB, PS; Methodology: SB, CB, PS; Writing-original draft preparation: CB; Writing-reviewing and editing: CB, SB, PS.

## Acknowledgements

The authors thank Nikita Woodhead for *Xenopus* care, Joanna Ward for general lab technical assistance, and Erin Damsteegt for assistance with *Xenopus* inductions.

## Literature cited

Bar-Klein, G., L. P. Cacheaux, L. Kamintsky, O. Prager, I. Weissberg et al., 2014 Losartan prevents acquired epilepsy via TGF-beta signaling suppression. Ann Neurol 75: 864–875.

Bell, M. R., J. A. Belarde, H. F. Johnson and C. D. Aizenman, 2011 A neuroprotective role for polyamines in a Xenopus tadpole model of epilepsy. Nat Neurosci 14: 505–512.

Brinkman, E. K., T. Chen, M. Amendola and B. van Steensel, 2014 Easy quantitative assessment of genome editing by sequence trace decomposition. Nucleic Acids Res 42: e168.

Camfield, C., and P. Camfield, 2008 Twenty years acer childhood-onset symptomatic generalized epilepsy the social outcome is usually dependency or death: a population-based study. Dev Med Child Neurol 50: 859–863.

Currie, S. P., D. Combes, N. W. Scott, J. Simmers and K. T. Sillar, 2016 A behaviorally related developmental switch in nitrergic modulation of locomotor rhythmogenesis in larval Xenopus tadpoles. J Neurophysiol 115: 1446–1457.

De Jesus Andino, F., L. Jones, S. B. Maggirwar and J. Robert, 2016 Frog Virus 3 dissemination in the brain of tadpoles, but not in adult Xenopus, involves blood brain barrier dysfunction. Sci Rep 6: 22508.

Demarest, S., J. Calhoun, K. Eschbach, H. C. Yu, D. Mirsky et al., 2022 Whole-exome sequencing and adrenocorJcotropic hormone therapy in individuals with infantile spasms. Dev Med Child Neurol 64: 633–640.

Doege, C., M. Luedde and K. Kostev, 2022 Association Between Angiotensin Receptor Blocker Therapy and Incidence of Epilepsy in Patients With Hypertension. JAMA Neurol 79: 1296–1302.

Du, C., F. Zheng and X. Wang, 2016 Exploring novel AEDs from drugs used for treatment of non-epileptic disorders. Expert Rev Neurother 16: 449–461.

Epi4K, P. Epilepsy Phenome/Genome, A. S. Allen, S. F. Berkovic, P. Cossette et al., 2013 De novo mutations in epileptic encephalopathies. Nature 501: 217–221.

Epi, K. C., 2016 De Novo Mutations in SLC1A2 and CACNA1A Are Important Causes of Epileptic Encephalopathies. Am J Hum Genet 99: 287–298.

Exner, C. R. T., and H. R. Willsey, 2021 Xenopus leads the way: Frogs as a pioneering model to understand the human brain. Genesis 59: e23405.

Fisher, M., C. James-Zorn, V. Ponferrada, A. J. Bell, N. Sundararaj et al., 2023 Xenbase: key features and resources of the Xenopus model organism knowledgebase. Genetics 224.

Fujimura, K., A. J. Guise, T. Nakayama, C. N. Schlaffner, A. Meziani et al., 2023 Integrative systems biology characterizes immune-mediated neurodevelopmental changes in murine Zika virus microcephaly. iScience 26: 106909.

Gorter, J. A., E. A. van Vliet and E. Aronica, 2015 Status epilepticus, blood-brain barrier disruption, inflammation, and epileptogenesis. Epilepsy Behav 49: 13–16.

Gross, R. A., 1992 A brief history of epilepsy and its therapy in the Western Hemisphere. Epilepsy Res 12: 65–74.

Hewapathirane, D. S., D. Dunfield, W. Yen, S. Chen and K. Haas, 2008 In vivo imaging of seizure activity in a novel developmental seizure model. Exp Neurol 211: 480–488.

Hong, S., H. JianCheng, W. JiaWen, Z. ShuQin, Z. GuiLian et al., 2019 Losartan inhibits development of spontaneous recurrent seizures by preventing astrocyte activation and aienuaJng blood-brain barrier permeability following pilocarpine-induced status epilepticus. Brain Res Bull 149: 251–259.

Jamieson, D., and A. Roberts, 2000 Responses of young Xenopus laevis tadpoles to light dimming: possible roles for the pineal eye. J Exp Biol 203: 1857–1867.

Keezer, M. R., G. S. Bell, A. Neligan, J. Novy and J. W. Sander, 2016 Cause of death and predictors of mortality in a community-based cohort of people with epilepsy. Neurology 86: 704–712.

Kimmel, C. B., J. Patterson and R. O. Kimmel, 1974 The development and behavioral characteristics of the startle response in the zebra fish. Dev Psychobiol 7: 47–60.

Klein, P., A. Friedman, M. Q. Hameed, R. M. Kaminski, G. Bar-Klein et al., 2020 Repurposed molecules for antiepileptogenesis: Missing an opportunity to prevent epilepsy? Epilepsia 61: 359–386.

Labun, K., T. G. Montague, J. A. Gagnon, S. B. Thyme and E. Valen, 2016 CHOPCHOP v2: a web tool for the next generation of CRISPR genome engineering. Nucleic Acids Res 44: W272–276.

Li, V. J., A. Schohl and E. S. Ruthazer, 2022 Topographic map formation and the effects of NMDA receptor blockade in the developing visual system. Proc Natl Acad Sci U S A 119.

Loscher, W., 2020 The holy grail of epilepsy prevention: Preclinical approaches to antiepileptogenic treatments. Neuropharmacology 167: 107605.

McTague, A., K. B. Howell, J. H. Cross, M. A. Kurian and I. E. Scheffer, 2016 The genetic landscape of the epileptic encephalopathies of infancy and childhood. Lancet Neurol 15: 304–316.

Mis, E. K., A. G. Sega, R. H. Signer, T. Cartwright, W. Ji et al., 2021 Expansion of NEUROD2 phenotypes to include developmental delay without seizures. Am J Med Genet A.

Molnar, Z., C. Metin, A. Stoykova, V. Tarabykin, D. J. Price et al., 2006 Comparative aspects of cerebral cortical development. Eur J Neurosci 23: 921–934.

Moreno, N., and A. Gonzalez, 2017 Pattern of Neurogenesis and Identification of Neuronal Progenitor Subtypes during Pallial Development in Xenopus laevis. Front Neuroanat 11: 24.

Naert, T., D. Tulkens, N. A. Edwards, M. Carron, N. I. Shaidani et al., 2020 Maximizing CRISPR/Cas9 phenotype penetrance applying predictive modeling of editing outcomes in Xenopus and zebrafish embryos. Sci Rep 10: 14662.

Nieuwkoop, P. D., and J. Faber, 1967 A normal table of Xenopus laevis (Daudin). Elsevier/North Holland, Amsterdam.

Oliver, K. L., I. E. Scheffer, M. F. Bennett, B. E. Grinton, M. Bahlo and S. F. Berkovic, 2023 Genes4Epilepsy: An epilepsy gene resource. Epilepsia n/a.

Olson, J. M., A. Asakura, L. Snider, R. Hawkes, A. Strand et al., 2001 NeuroD2 is necessary for development and survival of central nervous system neurons. Dev Biol 234: 174–187.

Panthi, S., P. A. Chapman, P. Szyszka and C. W. Beck, 2023 Characterisation and automated quantification of induced seizure-related behaviours in Xenopus laevis tadpoles. J Neurochem.

Paridaen, J. T., and W. B. Huttner, 2014 Neurogenesis during development of the vertebrate central nervous system. EMBO Rep 15: 351–364.

Pawlik, M. J., B. Miziak, A. Walczak, A. Konarzewska, M. Chroscinska-Krawczyk et al., 2021 Selected Molecular Targets for Antiepileptogenesis. Int J Mol Sci 22.

Pereira, M. G., C. Becari, J. A. Oliveira, M. C. Salgado, N. Garcia-Cairasco and C. M. Costa-Neto, 2010 Inhibition of the renin-angiotensin system prevents seizures in a rat model of epilepsy. Clin Sci (Lond) 119: 477–482.

Poke, G., J. Stanley, I. E. Scheffer and L. G. Sadleir, 2023 Epidemiology of Developmental and Epileptic Encephalopathy and of Intellectual Disability and Epilepsy in Children. Neurology 100: e1363–e1375.

Pratt, K. G., and A. S. Khakhalin, 2013 Modeling human neurodevelopmental disorders in the Xenopus tadpole: from mechanisms to therapeutic targets. Dis Model Mech 6: 1057–1065.

Radu, B. M., F. B. Epureanu, M. Radu, P. F. Fabene and G. Bertini, 2017 Nonsteroidal anJ-inflammatory drugs in clinical and experimental epilepsy. Epilepsy Res 131: 15–27.

Reyes-Garcia, S. Z., C. A. Scorza, N. N. Ortiz-Villatoro and E. A. Cavalheiro, 2019 Losartan fails to suppress epileptiform activity in brain slices from resected tissues of patients with drug resistant epilepsy. J Neurol Sci 397: 169–171.

Runge, K., R. Mathieu, S. Bugeon, S. Lafi, C. Beurrier et al., 2021 Disruption of NEUROD2 causes a neurodevelopmental syndrome with autistic features via cell-autonomous defects in forebrain glutamatergic neurons. Mol Psychiatry 26: 6125–6148.

Sakpichaisakul, K., R. Boonkrongsak, P. Lertbutsayanukul, N. Iemwimangsa, S. Klumsathian et al., 2022 Epileptic spasms related to neuronal differentiation factor 2 (NEUROD2) mutation respond to combined vigabatrin and high dose prednisolone therapy. BMC Neurol 22: 461.

Scheffer, I. E., S. Berkovic, G. Capovilla, M. B. Connolly, J. French et al., 2017 ILAE classification of the epilepsies: Position paper of the ILAE Commission for Classification and Terminology. Epilepsia 58: 512–521.

Scheffer, I. E., and J. Liao, 2020 Deciphering the concepts behind “Epileptic encephalopathy” and “Developmental and epileptic encephalopathy". Eur J Paediatr Neurol 24: 11–14.

Schindelin, J., I. Arganda-Carreras, E. Frise, V. Kaynig, M. Longair et al., 2012 Fiji: an open-source plaÜorm for biological-image analysis. Nature Methods 9: 676–682.

Sega, A. G., E. K. Mis, K. Lindstrom, S. Mercimek-Andrews, W. Ji et al., 2019 De novo pathogenic variants in neuronal differentiation factor 2 (NEUROD2) cause a form of early infantile epileptic encephalopathy. J Med Genet 56: 113–122.

Session, A. M., Y. Uno, T. Kwon, J. A. Chapman, A. Toyoda et al., 2016 Genome evolution in the allotetraploid frog Xenopus laevis. Nature 538: 336–343.

Shen, M. W., M. Arbab, J. Y. Hsu, D. Worstell, S. J. Culbertson et al., 2018 Predictable and precise template-free CRISPR editing of pathogenic variants. Nature 563: 646–651.

Sillar, K. T., 2009 Mauthner cells. Curr Biol 19: R353–355.

Sillar, K. T., and R. M. Robertson, 2009 Thermal activation of escape swimming in post-hatching Xenopus laevis frog larvae. J Exp Biol 212: 2356–2364.

Spruiell Eldridge, S. L., J. F. K. Teetsel, R. A. Torres, C. H. Ulrich, V. V. Shah et al., 2022 A Focal Impact Model of Traumatic Brain Injury in Xenopus Tadpoles Reveals Behavioral Alterations, Neuroinflammation, and an Astroglial Response. Int J Mol Sci 23.

Ta, A. C., L. C. Huang, C. R. McKeown, J. E. Bestman, K. Van Keuren-Jensen and H. T. Cline, 2022 Temporal and spatial transcriptomic dynamics across brain development in Xenopus laevis tadpoles. G3 (Bethesda) 12.

Tchekalarova, J. D., N. Ivanova, D. Atanasova, D. M. Pechlivanova, N. Lazarov et al., 2016 Long-Term Treatment with Losartan Attenuates Seizure Activity and Neuronal Damage Without Affecting Behavioral Changes in a Model of Co-morbid Hypertension and Epilepsy. Cell Mol Neurobiol 36: 927–941.

Truszkowski, T. L. S., E. J. James, M. Hasan, T. J. Wishard, Z. Liu et al., 2016 Fragile X mental retardation protein knockdown in the developing Xenopus tadpole optic tectum results in enhanced feedforward inhibition and behavioral deficits. Neural Development 11: 14.

Willsey, H. R., C. R. T. Exner, Y. Xu, A. Everitt, N. Sun et al., 2021 Parallel in vivo analysis of large-effect autism genes implicates cortical neurogenesis and estrogen in risk and resilience. Neuron 109: 1409.

Willsey, H. R., Y. Xu, A. Everitt, J. Dea, C. R. T. Exner et al., 2020 The neurodevelopmental disorder risk gene DYRK1A is required for ciliogenesis and control of brain size in Xenopus embryos. Development 147.

Yoo, Y., J. Jung, Y. N. Lee, Y. Lee, H. Cho et al., 2017 GABBR2 Mutations Determine Phenotype in Rei Syndrome and Epileptic Encephalopathy. Annals of Neurology 82: 466–478.

Zarei, K., K. L. Elliott, S. Zarei, B. Fritzsch and J. H. J. Buchholz, 2017 A method for detailed movement pattern analysis of tadpole startle response. J Exp Anal Behav 108: 113–124.

